# Revealing the human mucinome

**DOI:** 10.1101/2021.01.27.428510

**Authors:** Stacy A. Malaker, Nicholas M. Riley, D. Judy Shon, Kayvon Pedram, Venkatesh Krishnan, Oliver Dorigo, Carolyn R. Bertozzi

**Author notes:** Correspondence should be addressed to S.A.M. and C.R.B. These authors contributed equally to the manuscript (Stacy A. Malaker, Nicholas M. Riley). **Author Contributions** S.A.M. and C.R.B. designed research; S.A.M., N.M.R., D.J.S., and K.P. performed research; V.K. and O.D. contributed human clinical samples; S.A.M., N.M.R., D.J.S., and K.P. analyzed data. S.A.M., N.M.R., and C.R.B. wrote the paper with input from all authors.

## Abstract

Mucin domains are densely O-glycosylated modular protein domains found in a wide variety of cell surface and secreted proteins. Mucin-domain glycoproteins are key players in a host of human diseases, especially cancer, but the scope of the mucinome remains poorly defined. Recently, we characterized a bacterial mucinase, StcE, and demonstrated that an inactive point mutant retains binding selectivity for mucins. In this work, we leveraged inactive StcE to selectively enrich and identify mucins from complex samples like cell lysate and crude ovarian cancer patient ascites fluid. Our enrichment strategy was further aided by an algorithm to assign confidence to mucin-domain glycoprotein identifications. This mucinomics platform facilitated detection of hundreds of glycopeptides from mucin domains and highly overlapping populations of mucin-domain glycoproteins from ovarian cancer patients. Ultimately, we demonstrate our mucinomics approach can reveal key molecular signatures of cancer from *in vitro* and *ex vivo* sources.

## Introduction

Mucin domains are modular protein domains that adopt rigid and extended bottle-brush like structures due to a high density of O-glycosylated serine and threonine residues^1–3^. Mucin-type O-glycans are characterized by an initiating α-N-acetylgalactosamine (α-GalNAc) monosaccharide that can be further elaborated into several core structures through complex regulation of glycosyltransferases^4,5^. As a result, mucin domains serve as highly heterogenous swaths of glycosylation that exert both biophysical and biochemical influence. For instance, this includes the ability to redistribute receptor molecules at the glycocalyx and to drive high avidity binding interactions^6–8^. In the canonical mucin (MUC) family, mucin domains occur as tandem repeats, creating heavily glycosylated superstructures. Canonical mucins are central to many functions in health and disease, and have long been associated with human cancers, e.g., MUC1 and MUC16 (also known as CA-125)^9–12^. Dysregulation of mucin domain expression and aberrant mucin domain glycosylation patterns have been implicated in disease pathologies, especially in tumor progression, where mucins modulate immune responses and also promote proliferation through biomechanical mechanisms^13–15^.

Mucin domains also exist in proteins outside of the 21 canonical mucins (**Fig 1A**). For example, CD43 on the surface of leukemia cells selectively interacts with the glyco-immune checkpoint receptor Siglec-7 through its N-terminal mucin domain^16^; mucin domain-containing splice variants of CD44 (CD44v) serve as cancer cell markers relative to the ubiquitously expressed standard isoform^17^; CD45 mucin domains act as suppressors of T-cell activation^18^; mucin domain O-glycosylation on PSGL-1 is required for leukocyte-endothelial interactions^19^; and aberrant regulation of mucin domains in podocalyxin and SynCAM1 are implicated in a variety of cancers^20,21^. In all of these cases, shared functional attributes of mucin domains impart structural and biophysical properties relevant to their biology. Thus, instead of the more traditional categorization of the glycoproteome into N- and O-glycoproteins (both of which are represented by mucin-domain glycoproteins), it is logical to parse the glycoproteome into the mucinome, a family of glycoproteins whose mucin domains make them functionally related. However, even as the tools to capture the broadly defined N- and O-glycoproteome continue to improve^22–31^, mucin domains remain enigmatic and difficult to characterize. As such, a comprehensive list of all proteins with a mucin domain does not exist. This lack of a well-defined mucinome leaves a critical blind spot in our ability to interrogate mucin domain functions across molecular biology.

**Figure 1.**
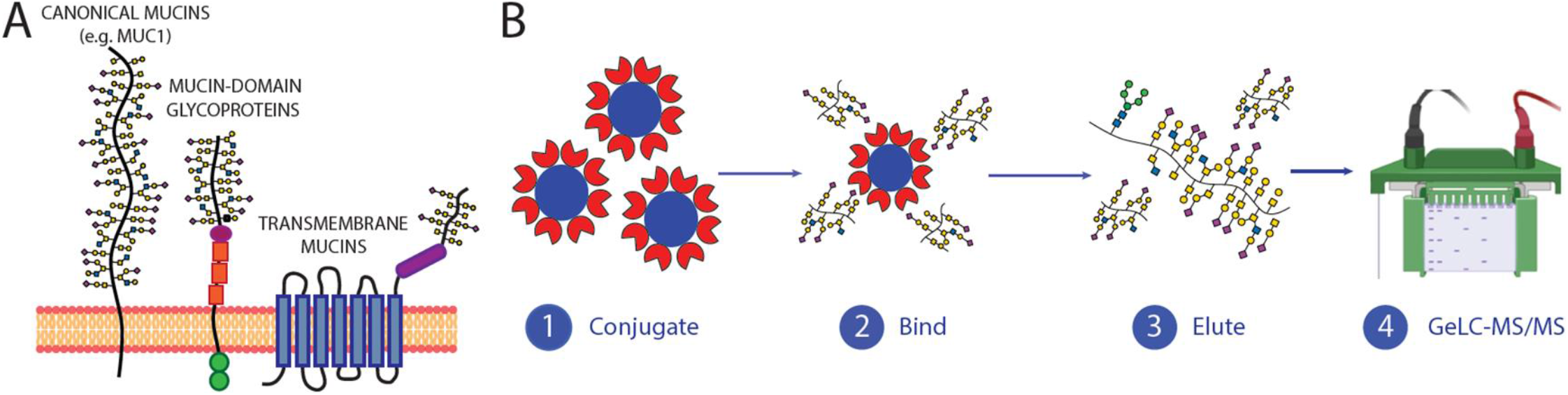
Mucinomics platform for enrichment of mucins in complex samples. **(A) The mucinome comprises a variety of proteins that have a densely glycosylated mucin domain.** Mucin domains are found in canonical mucins, mucin-domain glycoproteins, and even multi-pass transmembrane proteins. **(B) Workflow for enrichment technique.** StcE^E447D^ beads were conjugated using reductive amidation to POROS-AL 20 beads, followed by capping in Tris-HCl (1). Complex samples (lysate, ascites) were added to the beads and allowed to bind overnight (2), washed, and eluted by boiling in protein loading buffer (3). Samples were fractionated via one-dimensional gel electrophoresis, and digested in-gel using trypsin (4).

Toward this goal, enzymes derived from microorganisms known to colonize mucosal environments have shown promise for developing tools specifically suited to characterize mucin-domain glycoproteins^32–38^. We recently characterized a panel of such enzymes, termed mucinases, and showed that each of them harbor unique peptide- and glycan-based cleavage motifs^39^. Using catalytic point mutants, we also demonstrated that select mucinases can retain binding specificity for mucin domains; these were then used as mucin-selective staining reagents for Western blots, immunohistochemistry, and flow cytometry^39^. One particular mucinase of interest is secreted protease of C1 esterase inhibitor (StcE) from enterohemorrhagic *Escherichia coli*, which recognizes mucin domains decorated with a variety of O-glycan modifications^40–43^. This gives StcE both the selectivity needed to specifically bind mucin domains and the breadth to bind diverse mucin domain subtypes that vary in glycosylation patterns – lending its distinction as a “pan-mucinase”. Indeed, StcE has shown great utility for selective release of mucin fragments from biological samples and for improving mass spectrometry (MS)-based analysis of mucin domains^40^.

We reasoned that the catalytically inactive point mutant of StcE (StcE^E447D^) could function as a universal mucin enrichment tool for mucin domain discovery, similar to how O-glycosidases and engineered sialidases can enrich broadly for O-glycosylated and sialylated glycoproteins, respectively^44,45^. Here we show that StcE^E447D^-conjugated beads selectively enrich mucin-domain glycoproteins from complex cancer cell lysates and from crude ovarian cancer patient ascites fluid. As part of this workflow, we developed a mucin candidacy algorithm to assign confidence scores to proteins that have a high likelihood of containing a mucin domain. Additionally, we detected hundreds of glycopeptides derived from mucin domains in the StcE^E447D^-enriched samples. Ultimately, we demonstrate that this mucinomics platform can define key molecular signatures of cancer in both *in vitro* and *ex vivo* systems and is a novel approach to unravel the role of mucin domains in health and disease.

## Results

### Mucin enrichment and definition strategy to describe the mucinome

Our previous work indicated that a catalytically inactive point mutant of StcE (StcE^E447D^) retains its binding specificity for mucin domains while leaving them intact for subsequent analysis^39,40^. Through a straightforward reductive amidation approach, we conjugated StcE^E447D^ to POROS-AL beads to generate a solid phase support material to use for enrichments^46^. To optimize our enrichment protocol, we added StcE^E447D^-conjugated beads to OVCAR3 supernatant followed by an anti-MUC16 Western blot for detection. We tuned several parameters of the enrichment, including binding time, bead-to-substrate ratio, wash buffers, and elution conditions (*see* Methods); a simplified protocol is detailed in **Fig. 1B**. With a suitable enrichment protocol defined, we scaled up the reaction for mass spectrometry by enriching 500 μg of HeLa cell lysate with 100 μL of pre-washed StcE^E447D^-conjugated beads. Bound proteins were eluted by boiling in protein loading buffer, elutions were separated by one-dimensional gel electrophoresis, and in-gel digestions were performed prior to quantitative shotgun proteomics (i.e, GeLC-MS/MS, *see* **Supplementary Fig. 1**).

To calculate the degree of enrichment provided by StcE^E447D^-conjugated beads, 30 μg of unenriched cell lysate was simultaneously prepared and analyzed alongside each elution. Significantly enriched proteins were determined by comparing area-under-the-curve-based label free quantitation (LFQ) values for proteins in the elution relative to lysate, with processing and calculations performed using MaxQuant and Perseus^47,48^. The volcano plot in **Fig. 2A** shows several known and canonical mucins enriched in the elution (right; green), as opposed to untreated lysate (left). In particular, MUC1, MUC13, and MUC16 were significantly enriched, as well as known mucin-domain glycoproteins CD55 (decay-accelerating factor, DAF) and syndecan-1 (SDC1).

**Figure 2.**
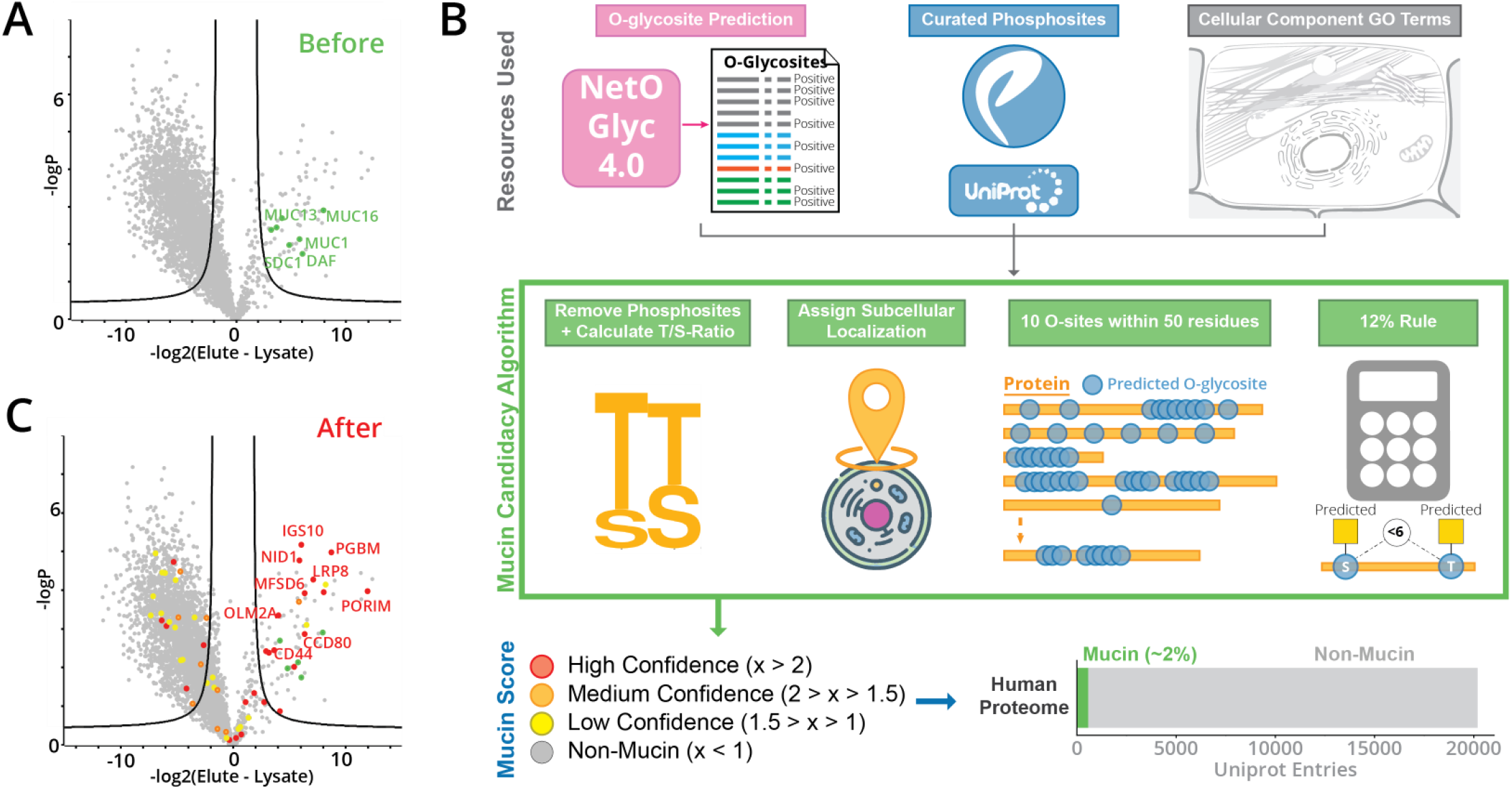
Mucin candidacy algorithm for confident assignment of mucin-domain glycoproteins. **(A) Known mucins in HeLa lysate enrichment.** HeLa lysate was subjected to the enrichment procedure described in Fig. 1 and known mucins (MUC1, MUC13, MUC16, DAF, and SDC1) were labeled. **(B) Mucin candidate annotation**. A mucin candidacy algorithm was created to assign Mucin Scores to indicate confidence that a given protein contains a mucin domain. First, predicted O-GalNAc sites were generated by the NetOGlyc4.0 tool, curated lists of phosphosites were downloaded from PhosphoSitePlus and Uniprot, and cellular localization GO terms were downloaded. The mucin candidacy algorithm then removed predicted O-GalNAc sites overlapping with known phosphosites, calculated the proportion of threonine to serine residues (T/S-ratio), evaluated protein subcellular localization, and checked for frequency and density of predicted O-GalNAc sites. These metrics were used to calculate a Mucin Score, which could then be used to evaluate mucinome enrichment. The entire human proteome was processed with the mucin candidacy algorithm; using manually curated benchmarks, approximately 330 proteins have mucin domains (~2% of human proteome). **(C) Mucinome of HeLa lysate.** The results in (A) were put through the mucin definition program, and mucins were labeled according to the Mucin Score. Red signified a score of >2 (high confidence), orange 2-1.5 (medium confidence), and yellow 1.5-1 (low confidence). Known mucins labeled in (A) are still labeled in green.

While these initial results were exciting, it quickly became clear that hand-curating proteins with known mucin domains would be untenable for the mucinome discovery platform. Not only is hand-curation low throughput, but it inherently misses proteins without known mucin domains. Instead, we developed a mucin candidacy algorithm to calculate which proteins have a high probability of bearing a mucin domain. As summarized graphically in **Fig. 2B**, our algorithm comprised several steps to assign a Mucin Score to every protein in the human proteome. Mucin candidacy algorithm processing is preceded by O-GalNAc glycosite prediction using the NetOGlyc4.0 tool, a support vector machine-based predictor developed using a map of ~3,000 O-glycosites from 600 O-glycoproteins that was generated through SimpleCell technology^49^. Predictions from NetOGlyc4.0 were then screened for known phosphosites annotated in Uniprot^50^ and PhosphoSitePlus^51^, and any overlap in phosphosites with predicted O-GalNAc sites resulted in removal of the predicted O-GalNAc site from consideration. This was a necessary step because NetOGlyc4.0 often predicted O-GalNAc sites in known phosphodomains of intracellular proteins, resulting in a high number of false positive mucin candidates after downstream processing. Note that O-GalNAc and phosphorylation sites are not known to have a high degree of overlap, as the former is generally extracellular whereas the latter is often intracellular.

Following O-GalNAc site prediction and phosphosite filtering, the algorithm asked four questions of each protein: (1) Was the protein predicted to be extracellular, secreted, and/or transmembrane?; (2) Were there at least 10 predicted O-glycosylation sites within a stretch of 50 residues?; (3) Was the distance between any given pair of O-glycosites less than 12% of the entire mucin domain (i.e., are glycosites <6 residues away from each other in a 50 residue sequence)?; and (4) Was the ratio of threonine to serine residues skewed toward threonine? Each of these benchmarks were determined through expert curation of known mucin sequences, which are further described in *Methods*. Using a point system based on the answers to these questions, the algorithm ultimately assigned a Mucin Score to each protein in the human proteome. By manually assessing outputs, we determined that a score of >2 was a high confidence mucin, between 2 and 1.5 was a medium confidence mucin, and between 1.5 and 1 was a low confidence mucin. Proteins with a score lower than 1 were considered non-mucins. Levels of confidence also capture the idea that a mucin domain may not be a binary concept; there may be gradients of O-glycosylation density and patterns that contribute to mucin-like attributes. See **Supplementary Dataset 1** for the mucin candidate algorithm output of the entire human proteome; approximately 330 proteins contain a mucin domain by our estimate (score > 1), encompassing all of the 21 canonical mucins, and comprising roughly 2% of the proteome (**Fig. 2B**). For comparison, proteases represent up to 2% of the human proteome; thus, mucin-domain glycoproteins could be much more common than previously thought^52^.

Using Mucin Scores to reannotate the dataset from Fig. 2B, we labeled high, medium, and low confidence mucins as red, orange, and yellow, respectively (**Fig. 2C**). The canonical and known mucins from Fig. 2B are still labeled in green. A large number of high confidence mucins are enriched in the StcE^E447D^ elution, which are labeled in red with gene names. Interestingly, some high confidence and a handful of medium to low confidence mucins are on the left side of the volcano plot, i.e., not enriched in the StcE^E447D^ elution. This could indicate (1) that StcE^E447D^ does not effectively enrich some mucin domains, (2) that mucin domains in these proteins are not heavily glycosylated in HeLa cells, or (3) that the mucin candidacy algorithm has some degree of error. Inherently, the mucin candidacy algorithm is an imperfect predictor of all mucin domains across the proteome. Indeed, no high efficacy mucin domain prediction algorithm exists, nor was that the focus of this work. Instead, our mucin candidacy algorithm indicates degrees of confidence for assigning proteins with a putative mucin domain that can be used to assess mucinome enrichment with StcE^E447D^-conjugated beads.

### Inactive mucinases enrich mucins from various cancer cell lines

Given that the HeLa lysate enrichment was successful, we decided to expand the approach to other cancer-associated cell lines, including SKBR3 (breast), OVCAR3 (ovarian), K562 (leukemia), and Capan2 (colorectal). The corresponding volcano plots are shown in **Fig. 3A-D**(*see* **Supplementary Dataset 2**). As before, red dots signified a score of >2 (high confidence), orange dots 2-1.5 (medium confidence), and yellow dots 1.5-1 (low confidence). Strongly enriched mucin-domain glycoproteins were labeled with their gene names. We also assigned a Boolean variable for each protein to denote its mucin domain status (“true” was assigned for Mucin Scores > 1) and performed Fisher’s exact tests in Perseus to derive enrichment scores for mucin status in elution and lysate proteomes. Here, higher values indicate a greater degree of enrichment, while values < 1 indicate depletion. For each cell line, enrichment factors > 14 were returned for proteins labeled as “true” (i.e., mucin-domain glycoproteins) in the elution compared to depletion factors between 0.5 and 0.7 for proteins labeled as “true” in the lysate (**Supplementary Fig. 2**). This indicates that mucin-domain glycoproteins are an enriched population in the elution and a depleted population in the lysate proteome.

**Figure 3.**
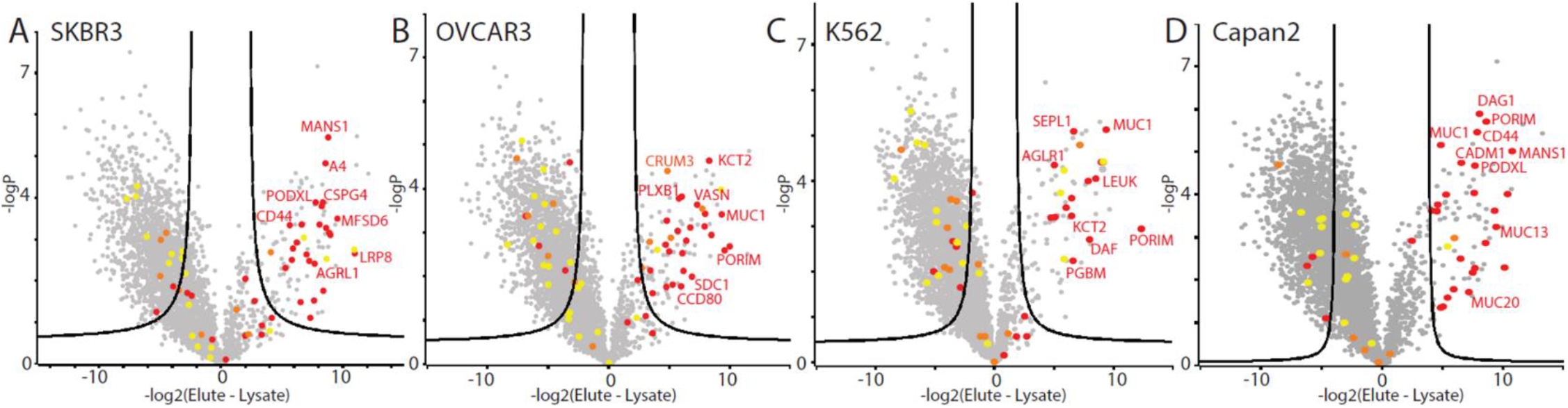
Mucinome of commonly used cancer cell lines. (A-D) Volcano plots representing four enrichment experiments from SKBR3 (A), OVCAR3 (B), K562 (C), and Capan2 (D) cell lines. Cell lysates were subjected to the workflow described in Fig. 1 and 2, scored with the mucin candidacy algorithm, and mucin-domain glycoproteins were labeled according to the Mucin Score. Red signified a score of >2 (high confidence), orange 2-1.5 (medium confidence), and yellow 1.5-1 (low confidence). Strongly enriched proteins were labeled with their gene names.

The Upset plot in **Fig. 4A** compares commonly observed mucin-domain glycoproteins across the cell lines. The total number of enriched mucins from each cell line is shown on the bottom left (blue horizontal bars). If a group of mucins was only seen in one cell line, only one gray dot is darkened; the number of proteins that are only seen in that cell line are shown in bar graph form above. For instance, 6 mucin-domain glycoproteins were only detected in the K562 cell line, whereas 5 mucin-domain glycoproteins were only detected in both the SKBR3 and OVCAR3 cell lines. Overlap between samples are shown by multiple darkened gray dots and a line connecting them. A total of six mucin-domain glycoproteins were seen in all five cell lines; these proteins are shown above the Upset plot. The putative mucin domain (orange, as calculated by the mucin candidacy algorithm), transmembrane domains (purple), and annotated N-glycan sites (green) are noted on each of the proteins. Several of the proteins were previously known to have a mucin domain, including: Mucin-1 (MUC1), Dystroglycan (DAG1), complement decay factor (CD55, DAF), and low-density lipoprotein receptor 8 (LRP8). However, we discovered that two of the overlapping proteins have previously undescribed mucin domains: major facilitator superfamily domain 6 (MSFD6) and Porimin (PORIM). The former is a multi-pass transmembrane protein that is implicated in antigen processing and presentation of exogenous peptide antigens via MHC class II, whereas porimin is involved in oncotic cell death characterized by vacuolization and increased membrane permeability. Neither of these proteins have annotated O-glycan sites and thus are “newly discovered” mucin-domain glycoproteins.

**Figure 4.**
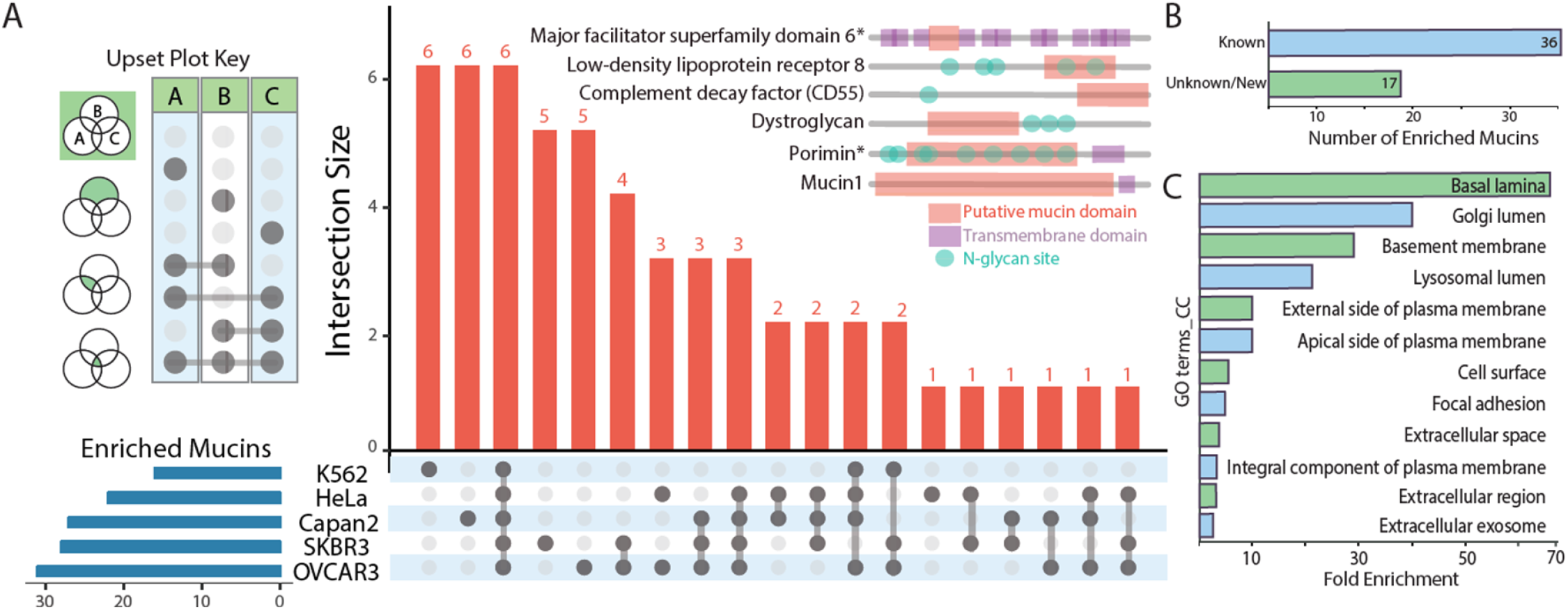
Analysis of mucin-domain glycoproteins from cell line enrichments. **(A) Upset plot comparing enriched mucin lists from five cell lines.** This graph shows intersections of mucin-domain glycoproteins from each cancer cell line. A key to reading the chart is shown at left. The six proteins found in all five samples are shown above the plot; putative mucin domain (orange), transmembrane domains (purple), and N-glycan sites (green) are annotated based on Uniprot assignments (transmembrane domain, N-glycan sites) and the mucin candidacy algorithm (mucin domain). **(B) Discovery of new mucin glycoproteins.** The total list of enriched mucin proteins was searched against the SimpleCell dataset^49^. If found in this dataset, it was considered a “known” mucin protein; if not, it was considered an unknown/new mucin glycoprotein. **(C) Mucin GO term enrichment.** Enriched cellular component (CC) GO_terms using DAVID^53,54^ are shown with fold enrichment indicated on the x-axis.

To better understand how many of the proteins contained previously undescribed mucin domains, we compared our dataset to the SimpleCell dataset from Clausen and colleagues^49^, which is the most comprehensive study on O-glycosites to date (albeit with truncated O-glycan species). In doing so, we found that approximately one-third (~31%, 17 of 55) were newly discovered mucin-domain glycoproteins (**Fig. 4B, Supplementary Table 1**). Of these proteins, perhaps the most surprising was adhesion G protein-coupled receptor L1 (ADGRL1), as GPCRs are generally not thought of as mucin-domain glycoproteins. This particular GPCR is implicated in both cell adhesion and signal transduction; future studies will be devoted to understanding the role of mucin domains in GPCR signaling. To broadly characterize features and functions of proteins present in our mucinome list, we performed GO term enrichment using DAVID^53,54^. Perhaps unsurprisingly, the most enriched cellular component (CC) GO terms were associated with membranes, cell surfaces, extracellular space, and cell adhesion (**Fig 4C**). The most common keywords were “heparan sulfate”, “proteoglycan”, “sulfation”, and “extracellular matrix” (**Supplementary Fig. 3**).

In a final extension of our mucinomics workflow, we performed an enrichment using a different mucinase. While StcE is a “pan-mucinase” (*vide infra*), we have characterized several other mucinases with varying proteolytic specificities^39^. BT4244 is a mucinase of particular interest from *Bacteroides thetaiotaomicron* that cleaves N-terminally to serine and threonine residues bearing truncated O-glycans, such as the cancer-associated T- and Tn-antigens (Gal-GalNAc and GalNAc, respectively). We reasoned that a point mutant of BT4244 (BT4244^E575A^) could also enrich mucin-domain glycoproteins bearing shortened O-glycan structures. Thus, we conjugated BT4244^E575A^ to beads and performed an analogous enrichment using HeLa lysate, with results shown in **Supplementary Fig. 4.** Only seven mucin-domain glycoproteins were significantly enriched in the elution, suggesting that not many mucins bear truncated O-glycans in HeLa cells. However, in the future, this proof-of-principle procedure could be used to enrich and identify cancer-associated glycoforms of mucins.

### Mucinomics platform allows for identification of ovarian cancer patient mucinome

Following the establishment of our mucin enrichment approach in cell lines, we next wanted to test the mucinomics platform on clinically relevant patient samples. Ovarian cancer ranks fifth in cancer deaths among women and is often diagnosed in stage III or IV, leading to a poor prognosis. This is due, in part, to the fact that the only clinically relevant biomarker is CA-125, a peptide epitope of MUC16, but the exact structural definition of this antigen continues to be elusive. Previously, we showed that StcE could digest MUC16 from crude ovarian cancer patient ascites fluid, leading us to reason that our enrichment technique could be used to selectively isolate MUC16 and other mucins from ascites fluid as a potential diagnostic strategy. As such, we performed the mucinomics enrichment with StcE^E447D^-beads on five de-identified patient samples (OC235, OC234, OC114, OC109, and OC107). As seen in **Fig. 5A-E**, a diverse array of mucins was significantly enriched in the elution (*see* **Supplementary Dataset 3**); in all but one of the experiments (OC114), MUC16 (denoted in purple) was significantly enriched. The enrichment in these experiments was even more successful than in the cell lines; in four out of five patient samples (excluding OC235), zero mucin-domain glycoproteins were “enriched” in the crude ascites fluid. For the full list of enriched mucin-domain glycoproteins, see **Supplemental Table 2.** Again, we compared our results to the SimpleCell dataset and found approximately one-third (~33%, 26 of 80) of the mucin candidates have previously unannotated mucin domains.

**Figure 5.**
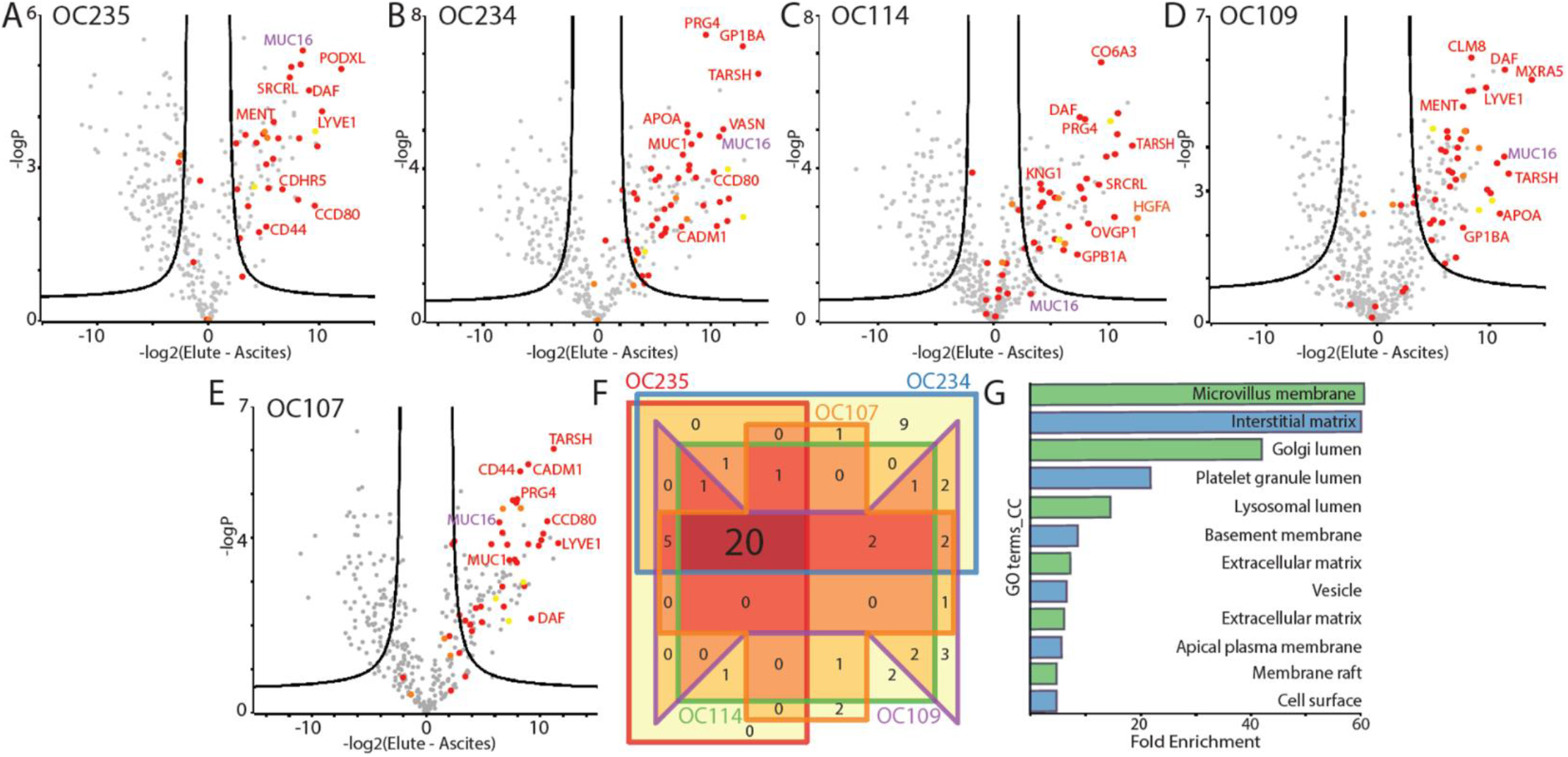
Mucinomics platform identifies ovarian cancer patient similarities. **(A-E) Volcano plots representing five enrichment experiments from crude ovarian cancer patient ascites fluid.** Ascites fluid samples were processed with the mucinomics workflow described in Fig. 1 and 2, including scoring with the mucin candidacy algorithm. Strongly enriched proteins are labeled with gene names. **(F) Venn diagram comparing mucinomics results from five cell lines.** Each sample is shown as a different color (red, orange, purple, green, and blue); 20 mucin proteins were enriched in all five samples. **(G) GO terms associated with mucin-domain glycoproteins.** Enriched CC GO terms using DAVID^53,54^ are shown with fold enrichment indicated on the x-axis

**Fig. 5F** compares overlap between the ascites samples with a Venn diagram of all enriched mucin proteins. Each sample is represented by a different color box, and the overlap between samples is given by a number within the boxes. Notably, 20 mucin-domain glycoproteins were enriched in all five samples, demonstrating substantial overlap between patients. The 20 overlapping proteins are listed in **Supplemental Table 3**. Again, as expected, the most enriched CC GO terms for the mucin-domain glycoproteins were associated with membranes, lumen, extracellular matrix, and the basement membrane (**Fig. 5G**). As before, the mucinome list contains some known mucin-domain glycoproteins, such as CD44, podocalyxin (PODXL), and agrin (AGRN). In addition, the list contains new mucins, such as thymosin beta-4 and Trem-like transcript 2 protein. Other proteins, like platelet glycoprotein Ib alpha chain (GP1BA) and lymphatic vessel endothelial hyaluronic acid receptor 1 (LYVE1), have previously been reported as “mucin-like”^55–57^, but do not have requisite evidence in the SimpleCell dataset, nor are any O-glycosites in the putative mucin domains reported on Uniprot. This further underscores the need for tools, like the strategy described here, to help define members of the mucinome. Additionally, we detected adhesion G protein-coupled receptor L1 (ADGRL1) as enriched in all five samples, further enforcing our conviction that this protein contains a mucin domain. While our patient cohort is currently too small to make any clinical claims, we believe that these overlapping mucins could be a better diagnostic and/or prognostic indicator for ovarian cancer. Future efforts will be devoted to expanding the study to a larger number of patients and comparing the results to patient outcomes, with the goal of developing a rapid mucin-fingerprinting approach using this mucinomics platform.

### StcE^E447D^-enrichment also captures O-glycopeptides from mucin domains

Characterization of intact O-glycopeptides was not an original goal of this platform, but StcE^E447D^-enrichment should function as a *de facto* glycopeptide enrichment by selecting for highly O-glycosylated mucins at the protein (i.e, pre-proteolysis) level. We observed a large number of spectra in our ascites enrichments bearing the “HexNAc fingerprint”, i.e., oxonium ions specific to glycopeptides, prompting us to search our data for intact glycopeptides. Generally, electron-driven dissociation is better suited for characterizing O-glycopeptides because it can provide O-glycosite localization.^58,59^ This is especially true for O-glycopeptides derived from mucins, which will likely have multiply glycosylated sequences^60–62^. Even so, collision-based fragmentation can still provide O-glycopeptide identifications that include peptide sequence and the total glycan mass modification, though details about number of glycans or glycosite positions are usually inaccessible. We collected only higher-energy collision dissociation (HCD) spectra through this study, limiting our ability to thoroughly characterize O-glycopeptides. Regardless, we searched our ascites data using O-Pair Search, a recently developed open-modification-centric glycoproteomic search algorithm that is particularly well-suited for the complex searches required of O-glycopeptide searches that consider large protein databases^63^. Even though we could not capitalize on the site-localization capabilities of O-Pair Search, we identified several hundred glycopeptides in both the enriched and crude ascites samples; the total list of all glycopeptides identified has been uploaded to the PRIDE database^64^.

Intriguingly, we discovered several O-glycopeptides on proteins that had previously uncharacterized mucin domains, as demonstrated in **Figure 6A**. Here, the putative mucin domain is indicated by an orange box, annotated N-glycan sites are shown with green dots, and approximate location of the O-glycopeptides detected are shown using red dots. These proteins do not have any annotated O-glycosites in the SimpleCell dataset or in Uniprot, thus these O-glycopeptides represent novel modifications on the mucin-domain glycoproteins. The presence of several identified O-glycopeptides in the regions assigned to be putative mucin domains by our mucin candidacy algorithm also strengthens our claim that the proteins do, in fact, have mucin domains. Additionally, we detected a large number of glycopeptides from MUC16, which is a key step toward better structural definition of this important cancer antigen.

**Figure 6.**
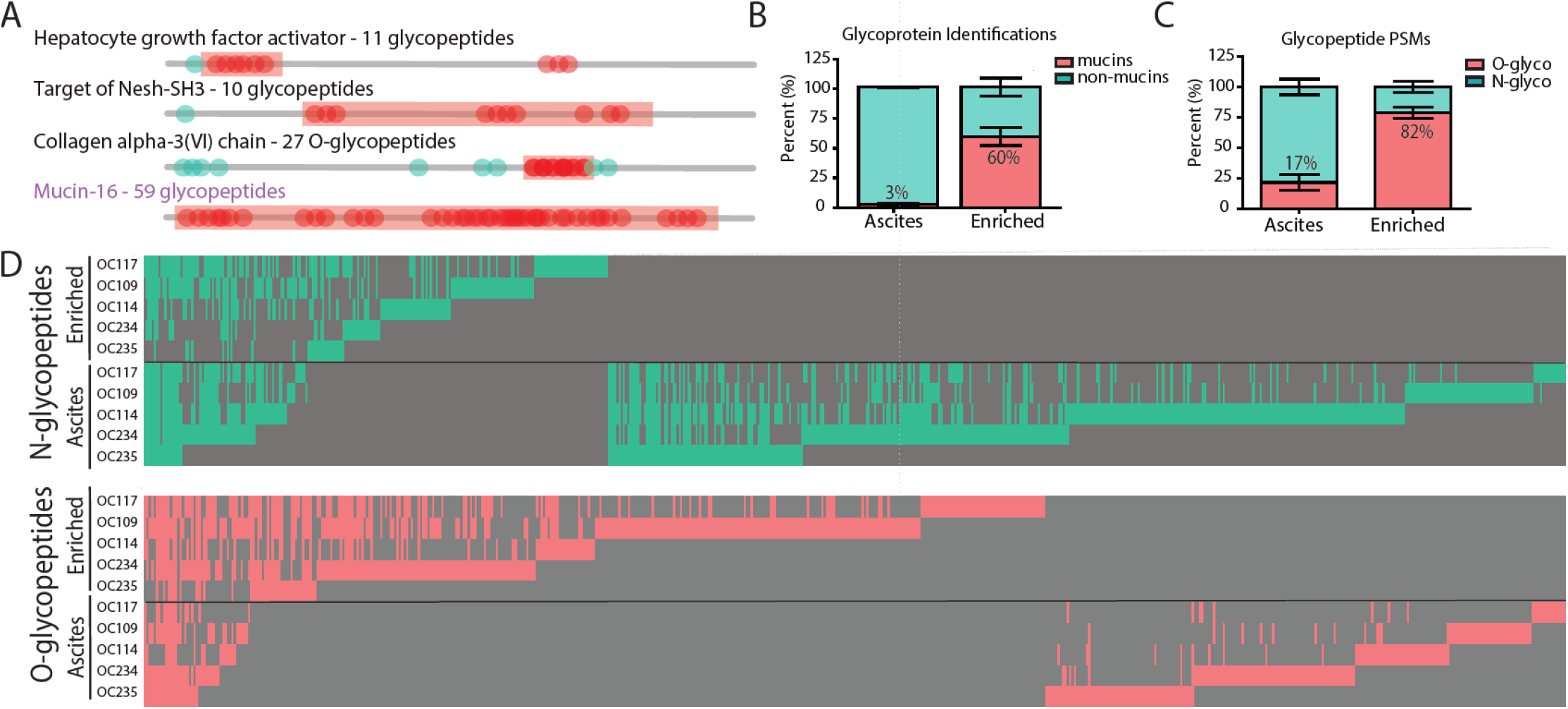
O-glycopeptides are highly abundant in the ascites enrichment. **(A) Mucin-domain glycoproteins harbor many O-glycosites**. O-glycopeptides identified by O-Pair Search are shown in red circles in approximate locations on the protein backbone; proteins were annotated based on Uniprot assignments (N-glycan sites, green) and the mucin candidacy algorithm (mucin domain, red). **(B) O-glycopeptide identifications map to mucin-domain glycoproteins identified with the mucin candidate algorithm**. Protein identifications from O-Pair glycopeptide searches were analyzed with the mucin candidacy algorithm, and this bar graph shows the percentage of protein identifications from the O-Pair Search that were non-mucin proteins (green) and mucin-domain glycoproteins (red). **(C) O-glycopeptide PSMs are higher in elution samples**. The total number of N- and O-glycopeptides were summed for each sample. The percentage of total glyco peptide spectral matches (PSMs) that were either N- or O-glycopeptides are shown in green and red, respectively. In both panels I and J, values represent the average of the 5 patient samples, and error bars show one standard deviation. **(D) O-glycopeptides are detected more often than N-glycopeptides in mucin-enriched ascites fluid.** Heat maps list all glycopeptides that were identified across all samples along the vertical axis, with samples along the top. Note, glycopeptide identifications along the vertical axis differ for the O- and N-glycopeptide heat maps. A glycopeptide detected in a given sample is indicated by color (green for N- and red for O-glycopeptides; top and bottom, respectively), while gray indicates the glycopeptide was not detected.

Next, we wanted to compare the glycoprotein sources of glycopeptides detected in the elution versus the crude cancer patient ascites fluid. As demonstrated in **Figure 6B,** only 3% of glycopeptides come from mucin-domain glycoprotein identifications in the unenriched ascites fluid, while 60% of glycopeptides from the elution come from mucins. Further, 82% of all glycopeptide identifications in the elution are O-glycopeptides (rather than N-glycopeptides), compared to only 17% in ascites fluid (**Fig. 6C**) This observation is also visualized in **Figure 6D**. Here, each vertical line represents one peptide and glycan combination. O-glycopeptides are on the bottom (pink), N-glycopeptides are on the top (green). The relative number of O-glycopeptides is much higher in the elution when compared to the ascites alone. The opposite is true for N-glycopeptides, which are detected more often in the ascites samples. Overall, these results provide further evidence that we can selectively enrich mucin-domain glycoproteins with a concomitant increase in O-glycopeptide identifications.

## Discussion

A rapidly developing breadth of tools continues to shed light on glycobiology, which is historically understudied relative to other biomolecules. Mucin-domain glycoproteins represent one particularly challenging subset of the glycoproteome that remains poorly defined. Though canonical mucins are recognized as important contributors to health and disease, a “parts list” for the mucinome, i.e., a complete list of mucin-domain glycoproteins, remains elusive, even though the mucinome is poised to address many open questions in glycobiology.

Here, we used a point mutant of our pan-mucinase, StcE^E447D^, along with a creative “mucin candidacy algorithm” to address this problem. While the enrichment feature of this approach appears robust, we note that the mucin candidacy algorithm is imperfect; yet, it serves a functional purpose for evaluating mucin-domain glycoprotein enrichments. Identification of mucin-domain glycoproteins in enriched in cell lysates rather than the elution could also indicate that certain mucin domains remain under-glycosylated depending on cellular state or cell type, meaning our mucinomics approach could be used to screen the mucin status of proteins under a variety of conditions. Our mucin candidacy algorithm could improve substantially from enhanced O-glycosite and mucin domain prediction tools. That said, prediction of mucin-type O-glycosites, much less mucin domains, remains challenging due to the complex regulation of O-glycosites by a poorly resolved family of glycosyltransferases. Future iterations could also explore other O-glycosite prediction algorithms beyond NetOGlyc4.0, such as ISOGlyP^65^.

Regardless, with this “mucinomics” platform, we enriched mucins from several cancer-associated cell lines and crude ovarian cancer patient ascites fluid. We demonstrated high mucin overlap between ovarian cancer patients, and the enrichment strategy allowed us to detect hundreds of glycopeptides from the mucin proteins, with a substantial increase in O- over N-glycopeptides. We also identified many proteins previously unknown to contain a mucin domain, thus demonstrating the utility of this technique in discovering new mucin-domain glycoproteins. Future efforts will be devoted to expanding our patient cohort in order to determine whether the ovarian cancer mucinome can be used as a diagnostic and/or prognostic indicator.

Though this work represents a significant step forward in understanding mucin domains, several open questions remain. To begin, mucin domains are known to regulate interactions at cellular peripheries via biophysical effects and cell-to-cell interactions. However, these roles are likely extremely dynamic, and may depend on various glycan structures (alone or in combination), expression of the mucin domain, and the overall cellular milieu. Further, the role of an individual mucin domain is unlikely to be identical across all of the mucin-domain glycoproteins. Thus, future studies should be devoted to understanding the role that discrete mucin domains are playing in cellular function. We predict that these mucin domains will fall into subgroups with categorical roles in health and disease.

Additionally, while we have identified a large number of mucin-domain glycoproteins from cell lines and ascites fluid, many other mucins are likely present on different cell types and in other indications. In particular, the immune cell mucinome is of incredible interest and may represent a class of new ‘checkpoint inhibitors’ with both glycan and peptide components to investigate^16^. Further, while we chose to focus our efforts on the cancer mucinome, several other disease mucinomes have yet to be studied. These mucinopathies include, but are not limited to, inflammatory bowel disease, cystic fibrosis, chronic obstructive pulmonary disease (COPD), Sjögren's syndrome, and dry mouth/eyes. Ultimately, we believe our mucinomics strategy will find utility in several settings, and will prove to be an invaluable tool for glycobiologists and biochemists alike.

## Methods

### Mucinase cloning, expression, and purification

StcE and BT4244 were expressed as previously described^39,40^. Briefly, Natalie Strynadka (University of British Columbia) kindly provided the plasmid pET28b-StcE-Δ35-NHis^43^. Robert Hirt (Newcastle University) kindly provided the plasmid pRSETA-BT4244^33^. pET28b-StcE^E447D^-Δ35-NHis and pRSETA-BT4244^E575A^ were generated using the Q5 Site-Directed Mutagenesis Kit (New England Biolabs) with primers listed in Table 1.

**Table 1.**
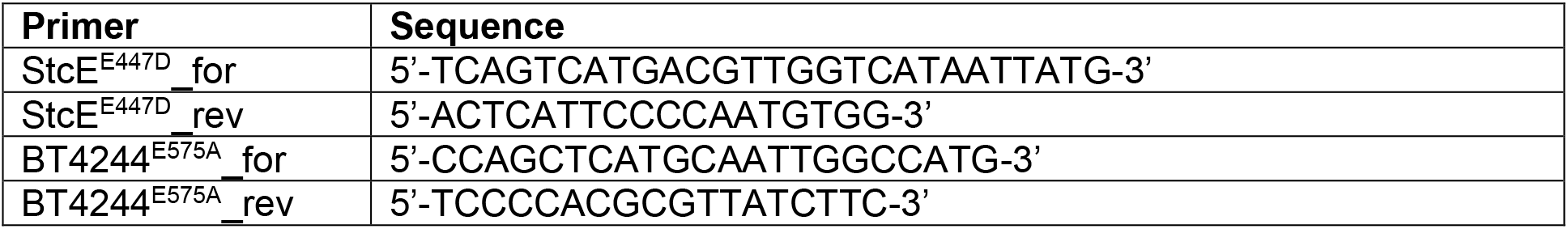
Cloning primers for inactive point mutants

StcE^E447D^ was expressed and purified as previously described^40^. BT4244^E575A^ was expressed in BL21(DE3) *E. coli* (New England Biolabs) grown in Luria broth (LB) with 100 μg/mL ampicillin at 37 °C, 225 rpm. The culture was induced at OD 0.6-0.8 with 1 mM IPTG and grown overnight at 20 °C. Cell pellets were lysed in xTractor buffer (Clontech) and lysates were applied to 1 mL HisTrap HP columns (Cytiva Life Sciences) using an ÄKTA Pure FPLC. Columns were washed with 50 column volumes of 20 mM Tris-HCl, 100 mM NaCl, 15 mM imidazole, pH 8, and elution was performed with a linear gradient to 150 mM imidazole. For BT4244, fractions containing pure protein were concentrated using Amicon Ultra 10 kDa MWCO filters (Millipore Sigma), dialyzed into PBS, pH 7.4, and stored at −80 °C. BT4244^E575A^ was further purified by size exclusion chromatography using a Superdex 200 Increase 10/300 GL column (Cytiva Life Sciences) in PBS, pH 7.4, and fractions containing pure protein were stored at −80 °C.

### Cell culture

Cells were maintained at 37 °C and 5% CO2. HeLa cells were cultured in DMEM supplemented with 10% fetal bovine serum (FBS) and 1% penicillin/streptomycin (P/S). Capan-2 cells were cultured in McCoy's 5a supplemented with 10% FBS and 1% P/S. K562 and SKBR3 cells were cultured in RPMI supplemented with 10% FBS and 1% P/S. OVCAR-3 cells were cultured in RPMI supplemented with 20% FBS, 0.01 mg/mL bovine insulin, and 1% P/S. To prepare lysate for pulldowns, cells plated in T75 flasks (Thermo Fisher Scientific) were grown until ~70% confluency, washed three times with DPBS, then lysed in 500 μL of RIPA buffer (Thermo Fisher Scientific) supplemented with EDTA-free protease inhibitor cocktail (Roche) and 0.1% benzonase (Millipore Sigma). Lysates were stored at −80 °C prior to pulldown.

### Bead derivatization

An aliquot containing approximately 2 mg of StcE^E447D^ (1 mL of 1.93 mg/mL) was added to 7-8 mg of POROS-AL beads, along with 1 μL of 80 mg/mL NaCNBH3. The reaction proceeded overnight, with shaking, at 4 °C. After conjugation, the beads were washed three times with 500 μL of ultrapure water, spinning at 8500 rpm for 5 min each time. To cap all excess aldehyde sites on the beads, 200 μL of Tris-HCl with 1 μL of 80 mg/mL NaCNBH3 was added to the beads. The reaction shook at room temperature for 2 h. Excess beads were stored at 4 °C for up to one month, and were washed before each enrichment. For BT4244^E575A^ conjugation, the enzyme concentration was 1.423 mg/mL, so 5.5 mg of POROS-AL beads was used. Otherwise, the conjugation and enrichment steps were identical.

### Enrichment of mucins from cell lysates and ascites fluid

Cell lysates were clarified by centrifuging for 20 min at 18,000 *x g*, and concentrations were determined using standard BCA assays. As per optimization experiments, the ideal ratio of lysate to beads (w/v) was determined to be 500 μg/100 μL, where 100 μL of the conjugated beads corresponded to 700 μg of beads in solution. The beads were pelleted at 8500 rpm for 5 min and the supernatant was removed. Then, 5 μL of 0.5 M EDTA and 500 μg of cell lysate was added to the beads and incubated at 4 °C overnight, with shaking. The reaction was performed six times, in tandem. After binding, the beads were spun at 8500 rpm for 5 min, and the supernatant was saved (“FT” or flow-through). Then, the beads were washed three times with 250 μL of PBS buffer containing 5 μL of 0.5 M EDTA. After the last wash, 32 μL of 4X protein loading buffer was added to the beads. For unenriched (control) samples, 30 μg of lysate was added to 10 μL of 4x protein loading buffer. All samples were then boiled at 95 °C for 5 min, spun for 2 min at 13,000 *x g*, and frozen for at least 1 h. The samples were then thawed and loaded onto 4-12% Criterion XT Bis-Tris precast gels (Bio-Rad), and run in 1x MOPS (Bio-Rad) for 90 minutes at 180 V. The total number of lanes for each experiment was 12, which included 6 control and 6 enriched lanes. After running, the lanes were stained using Bulldog Bio SafeStain and destained in ultrapure water. Eight bands were cut from each lane, giving a total of 96 slices per enrichment. The slices were frozen overnight at −80 °C.

For optimization and proof-of-principle purposes, only one replicate was performed, and all steps were run on a gel (FT, 3x washes, elution). Afterward, an anti-MUC16 Western blot was performed using anti-MUC16 antibody [X75] (Abcam) and IRDye® 800CW Goat anti-Mouse IgG (LI-COR Biosciences) according to manufacturer recommendations. Images (total protein, Western blot) were generated using an Odyssey CLx Near-Infrared Fluorescence Imaging System (LI-COR Biosciences).

Ascites fluid was obtained from O.D. and V.K. and was de-identified prior to our handling. Samples were selected based on their grouping in ref 28, as well as the amount of ascites available for the enrichment. In optimization experiments, the ideal ratio of lysate to beads (v/v) was determined to be 100 μL/100 μL, where 100 μL of the conjugated beads corresponded to 700 μg of beads in solution. Ascites was centrifuged at 4 °C at 18,000 *x g* for 20 min, and samples were removed from the supernatant. For control experiments, 6 μL of ascites was removed per lane for a total of 36 μL. Otherwise, the procedure was the same as above.

### In-gel digest and C18 clean-up for mass spectrometry

All slices were thawed in 200 μL of ultrapure water (Pierce), followed by a rinse with 200 μL of acetonitrile (ACN, Fisher). Fresh 50 mM ammonium bicarbonate (“AmBic”) was made, and samples were rinsed in 200 μL of AmBic for 20 min at RT. Afterward, samples were reduced using 5 mM dithiothreitol (DTT, Sigma) in AmBic for 35 min at 65 °C, with shaking, followed by alkylation using 50 mM iodoacetamide (IAA, Sigma) in AmBic for 30 min at RT, in the dark. Then, slices were rinsed once in AmBic, followed by two washes in fresh 50:50 AmBic:ACN for 10 min each. Slices were then dried in a vacuum concentrator, and rehydrated with 0.1 μg of trypsin in 200 μL of AmBic, and reacted overnight at 37 °C. The following day, samples were acidified with 2 μL of formic acid (FA, Thermo) and held at 37 °C for 45 min. The supernatant was discarded and 100 μL of 0.1% FA in 70% ACN was added to the slices for 30 min at 37 °C. The elution was collected, and the step was repeated once. The elution of adjacent slices was combined, for a total of 48 samples per enrichment. The resultant elution (400 μL) was dried in a vacuum concentrator).

All samples were subjected to desalting with a 96-well HyperSep C18 plate (Thermo). For all steps, solvent was added to the plate and centrifuged at 3000 rpm in a Sorvall Legend RT. To begin, wells were wet with 150 μL of ACN followed by equilibration with 150 μL of 0.1% FA in ultrapure water (“solvent A”). Samples were reconstituted in 150 μL of solvent A and added to the plate three times. The wells were then washed three times with 150 μL of solvent A, followed by elution three times using 100 μL of 0.1% FA in 80% ACN (“solvent B”). The combined elution for each sample (48), totaling 300 μL, was taken to dryness in a vacuum concentrator. All samples were reconstituted in 7 μL of solvent A.

### Mass spectrometry

Samples were analyzed by online nanoflow LC-MS/MS using an Orbitrap Fusion Tribrid mass spectrometer (Thermo Fisher Scientific) coupled to a Dionex Ultimate 3000 HPLC (Thermo Fisher Scientific). A portion of the sample was loaded via autosampler isocratically onto a C18 nano pre-column using 0.1% formic acid in water (“Solvent A”). For all cell lysate samples and enriched ascites fluid, 6.5 μL of sample was injected onto the column; for unenriched ascites fluid, 0.5-6.5 μL of sample was loaded onto the column as determined by peptide BCA. For pre-concentration and desalting, the column was washed with 2% ACN and 0.1% formic acid in water (“loading pump solvent”). Subsequently, the C18 nano pre-column was switched in line with the C18 nano separation column (75 μm × 250 mm EASYSpray containing 2 μm C18 beads) for gradient elution. The column was held at 45 °C using a column heater in the EASY-Spray ionization source (Thermo Fisher Scientific). The samples were eluted at a constant flow rate of 0.3 μL/min using a 90 min gradient. The gradient profile was as follows (min:% solvent B, 2% formic acid in acetonitrile) 0:3, 3:5, 93:25, 103:35, 104:90, 109:90, 110:3, 140:3. The instrument method used an MS1 resolution of 60,000 at FWHM 400 m/z, an AGC target of 3e5, and a mass range from 350 to 1,500 m/z. Dynamic exclusion was enabled with a repeat count of 3, repeat duration of 10 s, exclusion duration of 10 s. Only charge states 2-6 were selected for fragmentation. MS2s were generated at top speed for 3 s. HCD was performed on all selected precursor masses with the following parameters: isolation window of 2 m/z, 30% collision energy, orbitrap detection (resolution of 30,000), and an AGC target of 1e4 ions.

### Mucin candidacy algorithm

To build the mucin candidacy algorithm, the entire human proteome was first downloaded from Uniprot (20,365 entries) and parsed into FASTA files containing 150 entries each (a total of 136 files). Each file was individually uploaded to the NetOGlyc4.0 Server (http://www.cbs.dtu.dk/services/NetOGlyc/) for O-glycosite prediction^49^. NetOglyc4.0 results were saved as .csv files for further processing. Cellular component (CC) GO terms for the human proteome were also downloaded from Uniprot, and phosphosite annotations were downloaded from Uniprot and PhosphoSitePlus^50,51^. Predictions from NetOGlyc4.0 were then screened for known phosphosites, and any overlap in phosphosites with predicted O-GalNAc sites resulted in removal of the predicted O-GalNAc site from consideration. To annotate proteins as extracellular, secreted, and/or transmembrane, cellular component localization terms from Uniprot were checked for each protein entry. A protein was annotated as “extracellular” if its CC GO terms contained the phrases “Cell Membrane”, “Cell membrane”, “pass membrane protein”, “Secreted”, “extracellular”, or “Extracellular”. Proteins also received the “extracellular” distinction if they contained GO accessions of 0005887, 0016021, or 0005576. Because many proteins have multiple locations, “extracellular” proteins were further denoted as “exclusively extracellular” if their GO term lists did NOT include “Mitochondrion”, “Cyto”, “cyto”, “Nucl”, or “cytoplasmic side”. Next, predicted O-glycosites were iterated over to determine if a given protein would pass our “mucin test”, which consisted of two calculations. First, we required a protein to have at least ten predicted O-glycosites within a 50-residue region. If a protein qualified for this benchmark, we applied our “12% rule” to determine the number of residues that separated any two given O-glycosites within this 50-residue region. The 12% rule applied to a 50-residue region meant that fewer than 6 residues could separate any given pair of O-glycosites. Both the “10 sites within 50 residues” metric and the “12% rule” were derived through hand annotation of known and thoroughly studied mucins mostly curated by the Mucin Biology Group (http://www.medkem.gu.se/mucinbiology/databases/db/Mucin-human-2015.htm)^66,67^. Although this could be considered both too stringent or too relaxed depending on perspective, empirical testing showed these rules (in conjunction with the other metrics discussed) to be reasonably reliable in properly annotating known mucin domains. Exploration of these “mucin test” metrics in particular is an interesting area for future studies looking to employ a mucin candidacy algorithm. Finally, a threonine to serine ratio (T/S-ratio) was calculated for the predicted O-glycosites, mainly as a metric to discriminate O-GalNAc sites (slight threonine preference) from phosphosites (slight serine preference) due to the proclivity of NetOGlyc4.0 to predict dense regions of O-GalNAc sites in what are actually intracellular phosphorylation domains. Note, these preferences are based on empirical observations. If the number of serines and threonines were both greater than zero, the T/S-ratio was calculated by taking the number of threonines and dividing by the number of serines. If the number of threonines was > 0 but the number of serines was 0, the T/S-ratio was assigned a value of 2. Otherwise, the T/S-ratio was set at 0. With all of these determinations made, we then generated a Mucin Score. First, an integer score was calculated. If a protein was annotated as “extracellular” and passed the “mucin test”, it received an integer score of 1, while proteins “exclusively extracellular” and passing the mucin test received an integer score of 2. Integer scores could then be augmented in two ways: a) if the predicted number of O-glycosites was greater than the number of annotated phosphosites, 1 point was added; and b) if the number of predicted O-glycosites was greater than five and the protein name contained “mucin”, “Mucin”, “proteoglycan”, or “Proteoglycan”, 3 points were added. Finally, the integer score was multiplied by the T/S-ratio to generate the Mucin Score. This process was completed for all proteins in the human proteome that had predictions returned from NetOGlyc4.0 (~20,191 entries). Mucin Scores were used to determine confidence that a protein contained a mucin domain, including valuations of high confidence (Mucin Score > 2), medium confidence (2 > Mucin Score > 1.5), low confidence (1.5 > Mucin Score > 1), and non-mucin (Mucin Score < 1). This annotation was determined manually, by assessing all of the factors above.

### Unmodified peptide MS data analysis (MaxQuant)

Raw data were processed using MaxQuant version 1.6.3.4, and tandem mass spectra were searched with the Andromeda search algorithm. Oxidation of methionine and protein N-terminal acetylation were specified as variable modifications, while carbamidomethylation of cysteine was set as a fixed modification. A precursor ion search tolerance of 20 ppm and a product ion mass tolerance of 0.3 Da were used for searches, and two missed cleavages were allowed for full trypsin specificity. Peptide spectral matches were made against a target-decoy human reference proteome database downloaded from Uniprot. Peptides were filtered to a 1% FDR and a 1% protein FDR was applied according to the target-decoy method. Proteins were identified and quantified using at least one peptide (razor + unique), where razor peptide is defined as a non-unique peptide assigned to the protein group with the most other peptides (Occam’s razor principle). Proteins were quantified and normalized using MaxLFQ^68^ with a label-free quantification (LFQ) minimum ratio count of 1. LFQ intensities were calculated using the match between runs feature, and MS/MS spectra were required for LFQ comparisons. For quantitative Article comparisons, protein intensity values were log2-transformed before further analysis, and missing values were imputed from a normal distribution with width 0.3 and downshift value of 1.8 (that is, default values) using the Perseus software suite^48^. A Boolean value “IsAMucin” was also appended to each protein, with the value set as true if the Mucin Score was greater than 1. Mucin Scores and IsAMucin were input manually into MQ ‘protein groups’ txt files for manipulation in Perseus. Significance testing was performed in Perseus using a two-tailed t-test with 250 randomizations, an FDR of 0.01, and an S0 value of 2 (all volcano plots), or in Microsoft Excel using a two-tailed t-test with heteroscedastic variance. Proteins were sorted by their Mucin Score and highlighted in red if the score was higher than 2 (“high probability mucin”), orange if between 2-1.5 (“medium probability mucin”, and yellow if between 1.5 and 1 (“low probability mucin”). Upset plots and the 5-sample Venn diagram (Fig. 4A and 5F, respectively) were generated using the Intervene Shiny app (https://intervene.shinyapps.io/intervene/)^69^. GO term enrichments were performed using DAVID^53,54^, with the human proteome as a background.

### Glycopeptide MS data analysis (O-Pair Search)

For glycopeptide analysis, samples were loaded into MetaMorpheus in groups of 8, related to one individual replicate (e.g. “Lysate 1” slice 1-8)^63,70^. The human proteome was loaded into the database (downloaded from Uniprot June, 2016), and a “Glyco” search task was selected. For each group of 8 raw files, an N- and an O-glyco search was performed separately. Parameters for the O-Glycopeptide Search were as follows: O-glycan database “Oglycan.gdb” (the default 12-glycan database), keep top 50 candidates, Dissociation type “HCD” and child scan “null”, 4 maximum Oglycan allowed, with OxoniumIonFit on. For the N-Glycopeptide Search, all parameters were the same except the “NGlycan182.gdb” database was used. For general peptide parameters, the following features were used: tryptic cleavage, maximum missed 2 cleavages, maximum 2 modifications per peptide, with a peptide length of 5-60. Precursor mass tolerance was set to 10 ppm, product mass tolerance at 20, with a minimum score allowed of 3. Finally, carbamidomethyl of Cys was set as a fixed modification, whereas oxidation of Met was set as a variable modification. All glycopeptide hits were filtered to have a Q value of less than 0.01 and all decoy hits were removed. In O-glycopeptide searches, any peptides that had the “N-glyco sequon” as “TRUE” were also removed. Bar graphs in Fig. 6A and 6B were made using PRISM, and the heat map in Fig. 6D was generated using Microsoft Excel.

## Supporting information

Dataset 1_Mucin Candidacy Output

Dataset 2_Cell line Perseus results

Dataset 3_Ascites Perseus results

## Acknowledgements

We thank Natalie Strynadka (University of British Columbia) and Robert Hirt (Newcastle University) for their gifts of the StcE and BT4244 expression plasmids, respectively. We also thank C.C. Angelakos for his assistance with statistical calculations, and Jessica Stark and Rishikesh Kulkarni for helpful discussions. This work was supported, in part, by National Cancer Institute Grant R01CA200423 (to C.R.B.). S.A.M. was supported by a National Institute of General Medical Sciences F32 Postdoctoral Fellowship (F32-GM126663-01). N.M.R. was funded through an NIH Predoctoral to Postdoctoral Transition Award (Grant K00 CA212454). D.J.S. was supported by a National Science Foundation Graduate Research Fellowship and Stanford Graduate Fellowship. K.P. was supported by a National Science Foundation Graduate Research Fellowship, a Stanford Graduate Fellowship, and the Stanford Chemistry, Engineering & Medicine for Human Health (ChEM-H) Chemistry/Biology Interface Predoctoral Training Program.

## References

1. Shurer, C. R. et al. Physical Principles of Membrane Shape Regulation by the Glycocalyx. Cell 177, 1757–1770.e21 (2019).

2. Wagner, C. E., Wheeler, K. M. & Ribbeck, K. Mucins and Their Role in Shaping the Functions of Mucus Barriers. Annu. Rev. Cell Dev. Biol. 34, 189–215 (2018).

3. Hansson, G. C. Mucins and the Microbiome. Annual Review of Biochemistry 89, 769–793 (2020).

4. Bennett, E. P. et al. Control of mucin-type O-glycosylation: A classification of the polypeptide GalNAc-transferase gene family. Glycobiology 22, 736–756 (2012).

5. Reily, C., Stewart, T. J., Renfrow, M. B. & Novak, J. Glycosylation in health and disease. Nature Reviews Nephrology 15, 346–366 (2019).

6. Möckl, L. The Emerging Role of the Mammalian Glycocalyx in Functional Membrane Organization and Immune System Regulation. Front. Cell Dev. Biol. 8, 253 (2020).

7. Kuo, J. C. H., Gandhi, J. G., Zia, R. N. & Paszek, M. J. Physical biology of the cancer cell glycocalyx. Nat. Phys. 14, 658–669 (2018).

8. Singh, P. K. & Hollingsworth, M. A. Cell surface-associated mucins in signal transduction. Trends in Cell Biology 16, 467–476 (2006).

9. Kufe, D. W. Mucins in cancer: Function, prognosis and therapy. Nature Reviews Cancer 9, 874–885 (2009).

10. Jonckheere, N. & Van Seuningen, I. The membrane-bound mucins: From cell signalling to transcriptional regulation and expression in epithelial cancers. Biochimie 92, 1–11 (2010).

11. Bhatia, R. et al. Cancer-associated mucins: role in immune modulation and metastasis. Cancer and Metastasis Reviews 38, 223–236 (2019).

12. Hollingsworth, M. A. & Swanson, B. J. Mucins in cancer: Protection and control of the cell surface. Nature Reviews Cancer 4, 45–60 (2004).

13. Paszek, M. J. et al. The cancer glycocalyx mechanically primes integrin-mediated growth and survival. Nature 511, 319–325 (2014).

14. Woods, E. C. et al. A bulky glycocalyx fosters metastasis formation by promoting g1 cell cycle progression. Elife 6, (2017).

15. Van Putten, J. P. M. & Strijbis, K. Transmembrane Mucins: Signaling Receptors at the Intersection of Inflammation and Cancer. Journal of Innate Immunity 9, 281–299 (2017).

16. Wisnovsky, S. et al. Genome-wide CRISPR screens reveal a specific ligand for the glycan-binding immune checkpoint receptor Siglec-7. Proc. Natl. Acad. Sci. 118, e2015024118 (2021).

17. Wang, L., Zuo, X., Xie, K. & Wei, D. The role of CD44 and cancer stem cells. in Methods in Molecular Biology 1692, 31–42 (Humana Press Inc., 2018).

18. Xu, Z. & Weiss, A. Negative regulation of CD45 by differential homodimerization of the alternatively spliced isoforms. Nat. Immunol. 3, 764–771 (2002).

19. Carlow, D. A. et al. PSGL-1 function in immunity and steady state homeostasis. Immunological Reviews 230, 75–96 (2009).

20. Canals Hernaez, D. et al. PODO447: A novel antibody to a tumor-restricted epitope on the cancer antigen podocalyxin. J. Immunother. Cancer 8, (2020).

21. Murakami, Y. Involvement of a cell adhesion molecule, TSLC1/IGSF4, in human oncogenesis. Cancer Science 96, 543–552 (2005).

22. Sun, S. et al. Comprehensive analysis of protein glycosylation by solid-phase extraction of N-linked glycans and glycosite-containing peptides. Nat. Biotechnol. 34, 84–88 (2016).

23. Suttapitugsakul, S., Sun, F. & Wu, R. Recent Advances in Glycoproteomic Analysis by Mass Spectrometry. Analytical Chemistry 92, 267–291 (2020).

24. Riley, N. M., Hebert, A. S., Westphall, M. S. & Coon, J. J. Capturing site-specific heterogeneity with large-scale N-glycoproteome analysis. Nat. Commun. 10, (2019).

25. Khatri, K. et al. Comparison of Collisional and Electron-Based Dissociation Modes for Middle-Down Analysis of Multiply Glycosylated Peptides. J. Am. Soc. Mass Spectrom. 29, 1075–1085 (2018).

26. Woo, C. M. et al. Development of IsoTaG, a Chemical Glycoproteomics Technique for Profiling Intact N- and O-Glycopeptides from Whole Cell Proteomes. J. Proteome Res. 16, 1706–1718 (2017).

27. Thomas, D. R. & Scott, N. E. Glycoproteomics: growing up fast. Current Opinion in Structural Biology 68, 18–25 (2021).

28. Chernykh, A., Kawahara, R. & Thaysen-Andersen, M. Towards structure-focused glycoproteomics. Biochem. Soc. Trans. (2021). doi:10.1042/bst20200222

29. Yang, W., Ao, M., Hu, Y., Li, Q. K. & Zhang, H. Mapping the O-glycoproteome using site-specific extraction of O-linked glycopeptides (EXoO). Mol. Syst. Biol. 14, (2018).

30. Yang, S. et al. Deciphering Protein O-Glycosylation: Solid-Phase Chemoenzymatic Cleavage and Enrichment. Anal. Chem. 90, 8261–8269 (2018).

31. Levery, S. B. et al. Advances in mass spectrometry driven O-glycoproteomics. Biochimica et Biophysica Acta - General Subjects 1850, 33–42 (2015).

32. Ayala-Lujan, J. L. et al. Broad Spectrum Activity of a Lectin-Like Bacterial Serine Protease Family on Human Leukocytes. PLoS One 9, e107920 (2014).

33. Nakjang, S., Ndeh, D. A., Wipat, A., Bolam, D. N. & Hirt, R. P. A Novel Extracellular Metallopeptidase Domain Shared by Animal Host-Associated Mutualistic and Pathogenic Microbes. PLoS One 7, e30287 (2012).

34. Noach, I. et al. Recognition of protein-linked glycans as a determinant of peptidase activity. Proc. Natl. Acad. Sci. U. S. A. 114, E679–E688 (2017).

35. Henderson, I. R., Czeczulin, J., Eslava, C., Noriega, F. & Nataro, J. P. Characterization of Pic, a secreted protease of Shigella flexneri and enteroaggregative Escherichia coli. Infect. Immun. 67, 5587–5596 (1999).

36. Govindarajan, B. et al. A metalloproteinase secreted by Streptococcus pneumoniae removes membrane mucin MUC16 from the epithelial glycocalyx barrier. PLoS One 7, (2012).

37. Derrien, M. et al. Modulation of mucosal immune response, tolerance, and proliferation in mice colonized by the mucin-degrader Akkermansia muciniphila. Front. Microbiol. 2, (2011).

38. Florencia Haurat, M. et al. The glycoprotease CpaA secreted by medically relevant acinetobacter species targets multiple O-linked host glycoproteins. MBio 11, 1–19 (2020).

39. Shon, D. J. et al. An enzymatic toolkit for selective proteolysis, detection, and visualization of mucin-domain glycoproteins. Proc. Natl. Acad. Sci. U. S. A. 117, 21299–21307 (2020).

40. Malaker, S. A. et al. The mucin-selective protease StcE enables molecular and functional analysis of human cancer-associated mucins. Proc. Natl. Acad. Sci. U. S. A. 116, 7278–7287 (2019).

41. Lathem, W. W. et al. StcE, a metalloprotease secreted by Escherichia coli O157:H7, specifically cleaves C1 esterase inhibitor. Mol. Microbiol. 45, 277–288 (2002).

42. Grys, T. E., Walters, L. L. & Welch, R. A. Characterization of the StcE protease activity of Escherichia coli O157:H7. J. Bacteriol. 188, 4646–4653 (2006).

43. Yu, A. C. Y., Worrall, L. J. & Strynadka, N. C. J. Structural insight into the bacterial mucinase StcE essential to adhesion and immune evasion during enterohemorrhagic E. coli infection. Structure 20, 707–717 (2012).

44. Woods, R. J. et al. Engineered High-Specificity Affinity Reagents for the Detection of Glycan Sialylation. FASEB J. 33, 801.2–801.2 (2019).

45. Riley, N. M., Bertozzi, C. R. & Pitteri, S. J. A Pragmatic Guide to Enrichment Strategies for Mass Spectrometry-based Glycoproteomics. Mol. Cell. Proteomics mcp.R120.002277 (2020). doi:10.1074/mcp.r120.002277

46. Malaker, S. A. et al. Identification and Characterization of Complex Glycosylated Peptides Presented by the MHC Class II Processing Pathway in Melanoma. J. Proteome Res. 16, 228–237 (2017).

47. Tyanova, S., Temu, T. & Cox, J. The MaxQuant computational platform for mass spectrometry-based shotgun proteomics. Nat. Protoc. 11, 2301–2319 (2016).

48. Tyanova, S. et al. The Perseus computational platform for comprehensive analysis of (prote)omics data. Nature Methods (2016). doi:10.1038/nmeth.3901

49. Steentoft, C. et al. Precision mapping of the human O-GalNAc glycoproteome through SimpleCell technology. EMBO J. 32, 1478–1488 (2013).

50. Bateman, A. UniProt: A worldwide hub of protein knowledge. Nucleic Acids Res. 47, D506–D515 (2019).

51. Hornbeck, P. V. et al. PhosphoSitePlus, 2014: Mutations, PTMs and recalibrations. Nucleic Acids Res. 43, D512–D520 (2015).

52. Vergnolle, N. Protease inhibition as new therapeutic strategy for GI diseases. Gut 65, 1215–1224 (2016).

53. Huang, D. W., Sherman, B. T. & Lempicki, R. A. Systematic and integrative analysis of large gene lists using DAVID bioinformatics resources. Nat. Protoc. 4, 44–57 (2009).

54. Huang, D. W., Sherman, B. T. & Lempicki, R. A. Bioinformatics enrichment tools: Paths toward the comprehensive functional analysis of large gene lists. Nucleic Acids Res. (2009). doi:10.1093/nar/gkn923

55. Narimatsu, Y. et al. An Atlas of Human Glycosylation Pathways Enables Display of the Human Glycome by Gene Engineered Cells. Mol. Cell 75, 394–407.e5 (2019).

56. Bensing, B. A. et al. Novel aspects of sialoglycan recognition by the Siglec-like domains of streptococcal SRR glycoproteins. Glycobiology 26, cww042 (2016).

57. Banerji, S. et al. LYVE-1, a new homologue of the CD44 glycoprotein, is a lymph-specific receptor for hyaluronan. J. Cell Biol. 144, 789–801 (1999).

58. Darula, Z. & Medzihradszky, K. F. Analysis of mammalian O-glycopeptides - We have made a good start, but there is a long way to go. Molecular and Cellular Proteomics 17, 2–17 (2018).

59. Windwarder, M. & Altmann, F. Site-specific analysis of the O-glycosylation of bovine fetuin by electron-transfer dissociation mass spectrometry. J. Proteomics 108, 258–68 (2014).

60. Pap, A., Klement, E., Hunyadi-Gulyas, E., Darula, Z. & Medzihradszky, K. F. Status Report on the High-Throughput Characterization of Complex Intact O-Glycopeptide Mixtures. J. Am. Soc. Mass Spectrom. 29, 1210–1220 (2018).

61. Khoo, K. H. Advances toward mapping the full extent of protein site-specific O-GalNAc glycosylation that better reflects underlying glycomic complexity. Current Opinion in Structural Biology 56, 146–154 (2019).

62. Riley, N. M., Malaker, S. A., Driessen, M. & Bertozzi, C. R. Optimal Dissociation Methods Differ for N- and O-glycopeptides. J. Proteome Res. (2020). doi:10.1021/acs.jproteome.0c00218

63. Lu, L., Riley, N. M., Shortreed, M. R., Bertozzi, C. R. & Smith, L. M. O-Pair Search with MetaMorpheus for O-glycopeptide characterization. Nat. Methods 17, 1133–1138 (2020).

64. Perez-Riverol, Y. et al. The PRIDE database and related tools and resources in 2019: improving support for quantification data. Nucleic Acids Res. 47, D442–D450 (2019).

65. Mohl, J. E., Gerken, T. A. & Leung, M.-Y. ISOGlyP: de novo prediction of isoform-specific mucin-type O-glycosylation. Glycobiology (2020). doi:10.1093/glycob/cwaa067

66. Lang, T. et al. Searching the Evolutionary Origin of Epithelial Mucus Protein Components - Mucins and FCGBP. Mol. Biol. Evol. 33, 1921–1936 (2016).

67. Stavenhagen, K. et al. N- and O-glycosylation Analysis of Human C1-inhibitor Reveals Extensive Mucin-type O-Glycosylation*□S. Mol. Cell. Proteomics 17, 1225–1238 (2018).

68. Cox, J. et al. Accurate proteome-wide label-free quantification by delayed normalization and maximal peptide ratio extraction, termed MaxLFQ. Mol. Cell. Proteomics 13, 2513–26 (2014).

69. Khan, A. & Mathelier, A. Intervene: A tool for intersection and visualization of multiple gene or genomic region sets. BMC Bioinformatics 18, 287 (2017).

70. Solntsev, S. K., Shortreed, M. R., Frey, B. L. & Smith, L. M. Enhanced Global Post-translational Modification Discovery with MetaMorpheus. J. Proteome Res. 17, 1844–1851 (2018).

