## Supplementary Data for "Revealing the human mucinome"

**This PDF file includes:**

Supplementary Tables 1 to 3
Supplementary Figures 1 to 4
References

**Other supplemental materials for this manuscript include the following:**

Dataset 1: Mucin candidate algorithm human proteome output
Dataset 2: MaxQuant and data processing results from all cell line enrichments
Dataset 3: MaxQuant and data processing results from all ascites enrichments

| Known | Unknown | Protein Names |
|---|---|---|
| O00468 | O94910 | Adhesion G protein-coupled receptor L1 |
| P05067 | Q9BY67 | Cell adhesion molecule 1 |
| P08174 | Q6UVK1 | Chondroitin sulfate proteoglycan 4 |
| P14543 | Q5VV43 | Dyslexia-associated protein KIAA0319 |
| P15941 | Q13261 | Interleukin-15 receptor subunit alpha |
| P16070 | P34810 | Macrosialin (CD68) |
| P18827 | Q6ZSS7 | Major facilitator superfamily domain-containing protein 6 |
| P98160 | Q9ULD2 | Microtubule-associated tumor suppressor 1 |
| Q14114 | Q8N387 | Mucin-15 |
| Q14118 | P98088 | Mucin-5AC |
| Q6UXD5 | Q68BL7 | Olfactomedin-like protein 2A |
| Q6WRI0 | P07359 | Platelet glycoprotein Ib alpha chain |
| Q76M96 | Q8N131 | Porimin |
| Q8NC54 | Q9BUF7 | Protein crumbs homolog 3 |
| Q8WXI7 | Q92954 | Proteoglycan 4 |
| Q9H3R2 | Q9Y289 | Sodium-dependent multivitamin transporter |
| Q9ULI3 | Q9HCM3 | UPF0606 protein KIAA1549 |
| O00592 | | |
| O95274 | | |
| P02751 | | |
| P43121 | | |
| Q6EMK4 | | |
| Q9H8J5 | | |
| Q9NQX5 | | |
| Q9NR99 | | |
| P39060 | | |
| Q6ZVL6 | | |
| Q8N307 | | |
| Q9HAR2 | | |
| O75056 | | |
| P08575 | | |
| P16150 | | |
| P28908 | | |
| Q14242 | | |
| Q99102 | | |
| Q9H1U4 | | |

**Supplementary Table 1. Known and new mucin-domain glycoproteins detected in cell line enrichments.** The total list of mucins detected in the cell line enrichments (duplicates have been removed) was compared to the SimpleCell dataset[1]. Proteins that were detected in SimpleCell data are listed under “known”, proteins that were not listed in this dataset are listed under “unknown” with the full protein name listed to the right.

| Known | New | Protein Name |
|---|---|---|
| O00592 | P58397 | A disintegrin and metalloproteinase with thrombospondin motifs 12 |
| Q4ZHG4 | O94910 | Adhesion G protein-coupled receptor L1 |
| Q76M96 | P16112 | Aggrecan core protein |
| P08174 | P08519 | Apolipoprotein(a) |
| Q8WXI7 | Q9HBB8 | Cadherin-related family member 5 |
| P15941 | Q9BY67 | Cell adhesion molecule 1 |
| Q9NR99 | Q6UVK1 | Chondroitin sulfate proteoglycan 4 |
| A1L4H1 | Q9UGN4 | CMRF35-like molecule 8 |
| Q9BUN1 | P25940 | Collagen alpha-3(V) chain |
| P16070 | P12111 | Collagen alpha-3(VI) chain |
| Q6EMK4 | Q9NPY3 | Complement component C1q receptor |
| Q9ULI3 | Q04756 | Hepatocyte growth factor activator |
| O00468 | Q8WWV6 | High affinity immunoglobulin alpha and immunoglobulin mu Fc receptor |
| P13611 | P01042 | Kininogen-1 |
| P05155 | Q8TF66 | Leucine-rich repeat-containing protein 15 |
| P12259 | Q9Y5Y7 | Lymphatic vessel endothelial hyaluronic acid receptor 1 |
| P18827 | Q6W4X9 | Mucin-6 |
| Q9HC84 | Q13201 | Multimerin-1 |
| Q9H1U4 | Q68BL7 | Olfactomedin-like protein 2A |
| Q6WRI0 | Q12889 | Oviduct-specific glycoprotein |
| Q14118 | Q9UKJ1 | Paired immunoglobulin-like type 2 receptor alpha |
| Q99102 | P07359 | Platelet glycoprotein Ib alpha chain |
| Q9UGM3 | Q92954 | Proteoglycan 4 |
| P08575 | Q7Z7G0 | Target of Nesh-SH3 |
| O95274 | Q95274 | Thymosin beta-4 |
| Q8N307 | Q96H15 | TIMD4 |
| | Q5T2D2 | Trem-like transcript 2 protein |

**Supplementary Table 2. Known and new mucin-domain glycoproteins detected in ascites fluid enrichments.** The total list of mucins detected in the ascites fluid enrichments (duplicates have been removed) was compared to the SimpleCell dataset[1]. Proteins that were detected in SimpleCell data are listed under “known”, proteins that were not listed in this dataset are listed under “unknown” with the full protein name listed to the right.

| Uniprot Number | Protein Name |
|---|---|
| P07359 | Platelet glycoprotein Ib alpha chain |
| Q9HBB8 | Cadherin-related family member 5 |
| Q9ULI3 | Protein HEG homolog 1 |
| Q7Z7G0 | Target of Nesh-SH3 |
| Q6EMK4 | Vasorin |
| P08519 | Apolipoprotein(a) |
| Q9BY67 | Cell adhesion molecule 1 |
| A1L4H1 | Soluble scavenger receptor cysteine-rich domain-containing protein SSC5D |
| Q9NR99 | Matrix-remodeling-associated protein 5 |
| O00468 | Agrin |
| Q76M96 | Coiled-coil domain-containing protein 80 |
| Q9Y5Y7 | Lymphatic vessel endothelial hyaluronic acid receptor 1 |
| Q9UGN4 | CMRF35-like molecule 8 |
| O00592 | Podocalyxin |
| Q04756 | Hepatocyte growth factor activator |
| P08174 | Complement decay-accelerating factor |
| Q4ZHG4 | Fibronectin type III domain-containing protein 1 |
| P25940 | Collagen alpha-3(V) chain |
| Q9BUN1 | Protein MENT |
| P16070 | CD44 antigen |

**Supplementary Table 3. Overlapping mucin-domain glycoproteins from five ascites enrichments.** The enriched mucins from five cancer patient ascites fluid samples were compared, and 20 proteins were found significantly enriched in all five samples. The Uniprot numbers are listed to the left and the protein names are listed on the right.

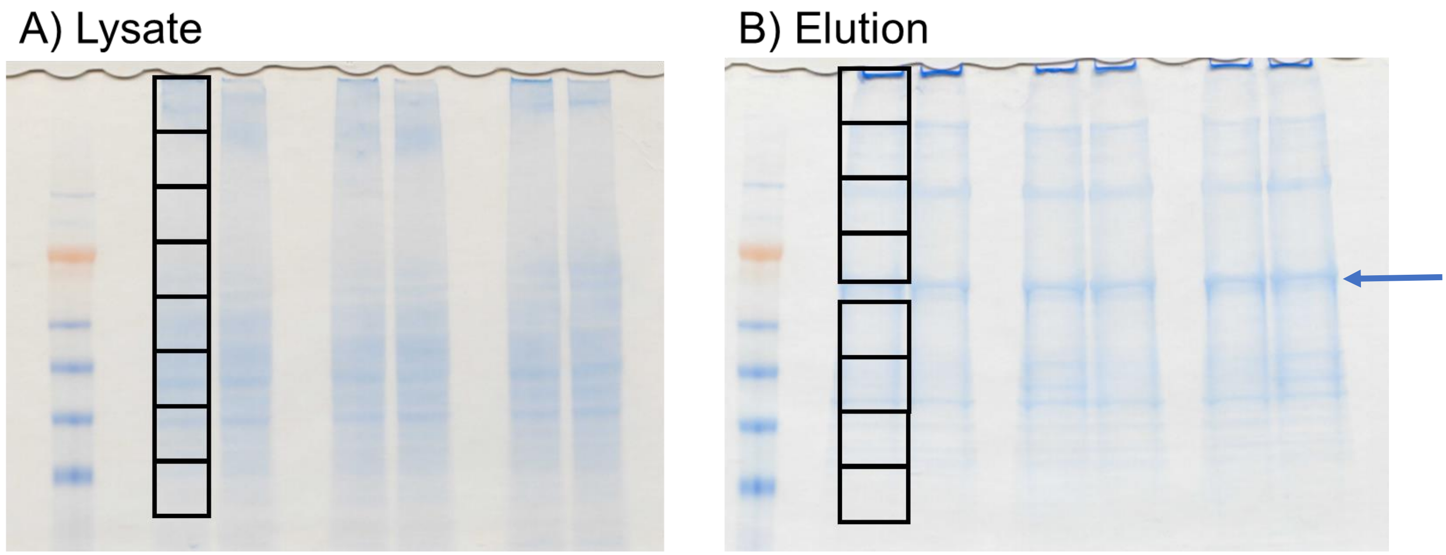


**Supplementary Figure 1. Example slices cut from lysate and elution for in-gel digests.** A) A total of 6 lanes were loaded with 6% of the enrichment input (~30 µg) and run on a 4-12% Bis-Tris gel for 1.5 h. Eight slices were cut from each lane; consecutive pairs of lanes were combined for a total of 3 replicates. B) A total of 6 lanes were loaded with the elution from 0.5 mg lysate in 100 µL beads (~200 µg StcE$^{E447D}$). Eight lanes were cut from each lane, avoiding the StcE$^{E447D}$ band (arrow). Consecutive pairs of lanes were combined for a total of 3 replicates.

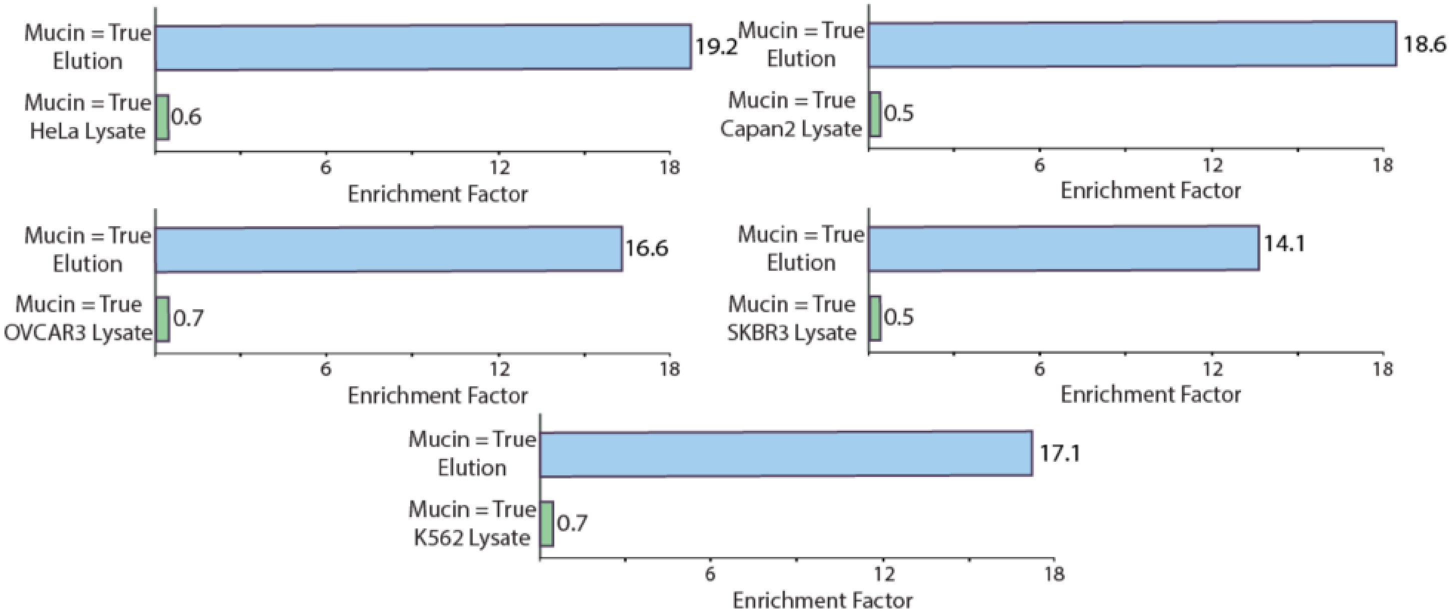


**Supplementary Figure 2. Enrichment factors for mucin-domain glycoproteins in cell line lysate and StcE$^{E447D}$ elution.** Enrichment factors were calculated using Fisher's exact tests in Perseus for all five cell line enrichments. In each case, the enrichment factor of "true", the Boolean variable assigned to proteins with a Mucin Score > 1, was >10x higher in the elution compared to the lysate alone. The average enrichment factor in the elution was 17.12, compared to an average of 0.6 in the lysate.

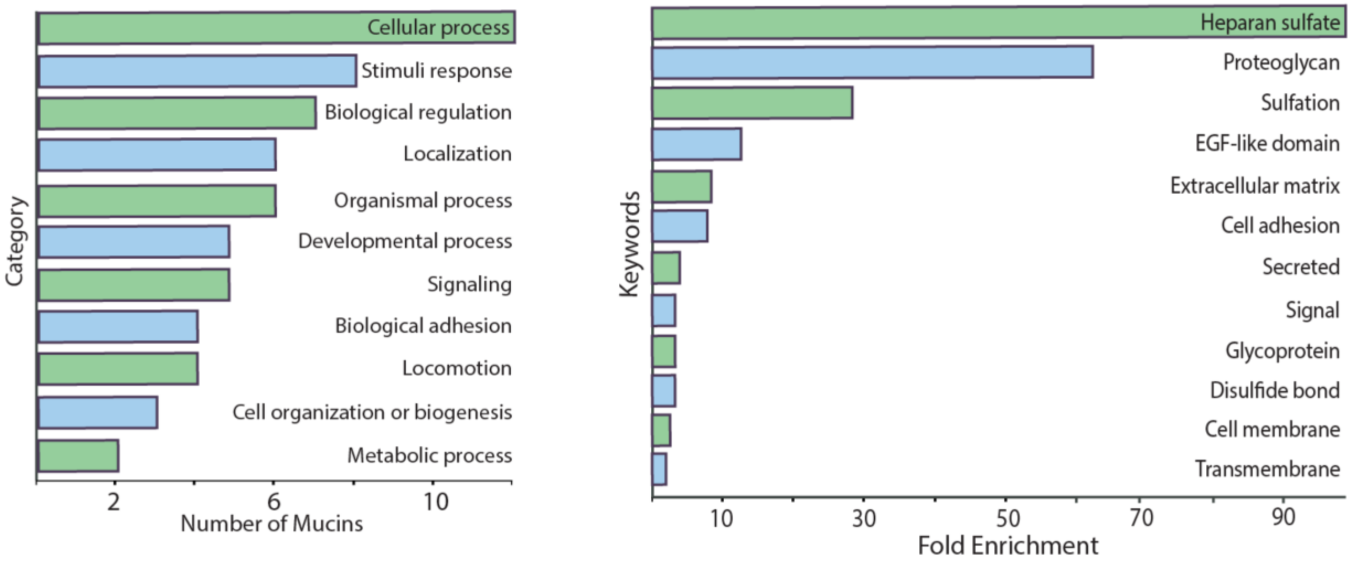


**Supplementary Figure 3. Enriched category and keywords of mucin-domain glycoproteins found in cell line enrichments.** The total list of enriched mucin proteins was input into DAVID, converted to gene names, and compared to the human genome as background. Enriched biological processes are shown on the left, enriched keywords are shown on the right.

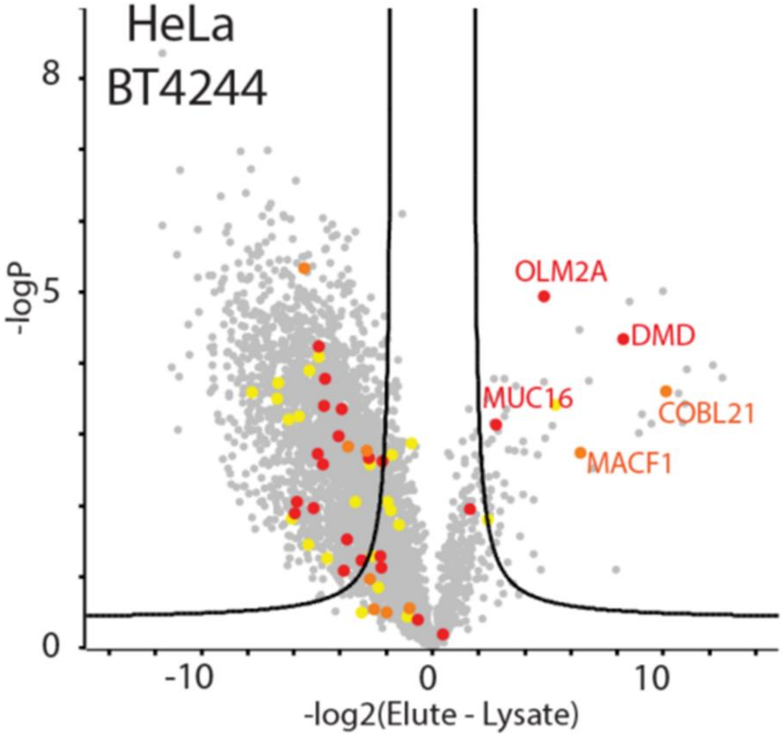


**Supplementary Figure 4. BT4244$^{E575A}$ enrichment of HeLa lysate.** BT4244$^{E575A}$ was conjugated to beads using reductive amidation and HeLa lysate was added to the beads. After binding, washing, and eluting, in-gel digest was performed on lysate alone and the elution of the enrichment. The samples were run on an Orbitrap Fusion Tribrid followed by a MaxQuant search. The data was processed using Perseus, and mucins were labeled according to the MucinScore. Red signified a score of >2 (high confidence), orange 2-1.5 (medium confidence), and yellow 1.5-1 (low confidence). Strongly enriched proteins are labeled with their gene names.

### References

1. Steentoft, C. *et al.* Precision mapping of the human O-GalNAc glycoproteome through SimpleCell technology. *EMBO J.* **32**, 1478–1488 (2013).
